# Spatially organized genomic and physiological heterogeneity of the olfactory bulb mitral cell layer

**DOI:** 10.1101/2020.01.13.903823

**Authors:** Daniel Paseltiner, Henry Loeffler, Alex Andonian, Abigail Leberman, Travis J. Gould, Jason B. Castro

**Affiliations:** Program in Neuroscience, Bates College, Lewiston ME 04240; Computer Science and Artificial Intelligence Laboratory (CSAIL), Massachusetts Institute of Technology, Cambridge, MA 02139; Physics Department, Bates College, Lewiston ME 04240

## Abstract

Sensory inputs to the dorsal vs ventral olfactory bulb derive from distinct receptor families, and drive distinct behaviors. To address whether second-order OB neurons and circuits exhibit matching heterogeneity for input-specific readout, we clustered spatial expression profiles of >2,000 genes from the mitral cell layer (MCL). We observed clear dorsal and ventral clusters, together with dorsoventral differences in mitral cell physiology. Bulbar circuits may therefore be tuned for zone-specific computation.

## Main

The main olfactory bulb receives topographically organized sensory inputs, partitioning this structure into columnar domains defined by individual glomeruli^1,2^, as well as larger dorsal and ventral glomerular zones defined by receptor clades^3–5^ (Fig. 1A). In genetic ablation studies of the OB’s zonal organization, it was found that the dorsal and ventral OB support innate vs. learned odor behaviors, respectively^6^. Consistent with this, dorsal and ventral portions of the bulb differ in their connectivity to downstream targets^7,8^ as well as their sources of centrifugal input^9^.

**Fig. 1.**
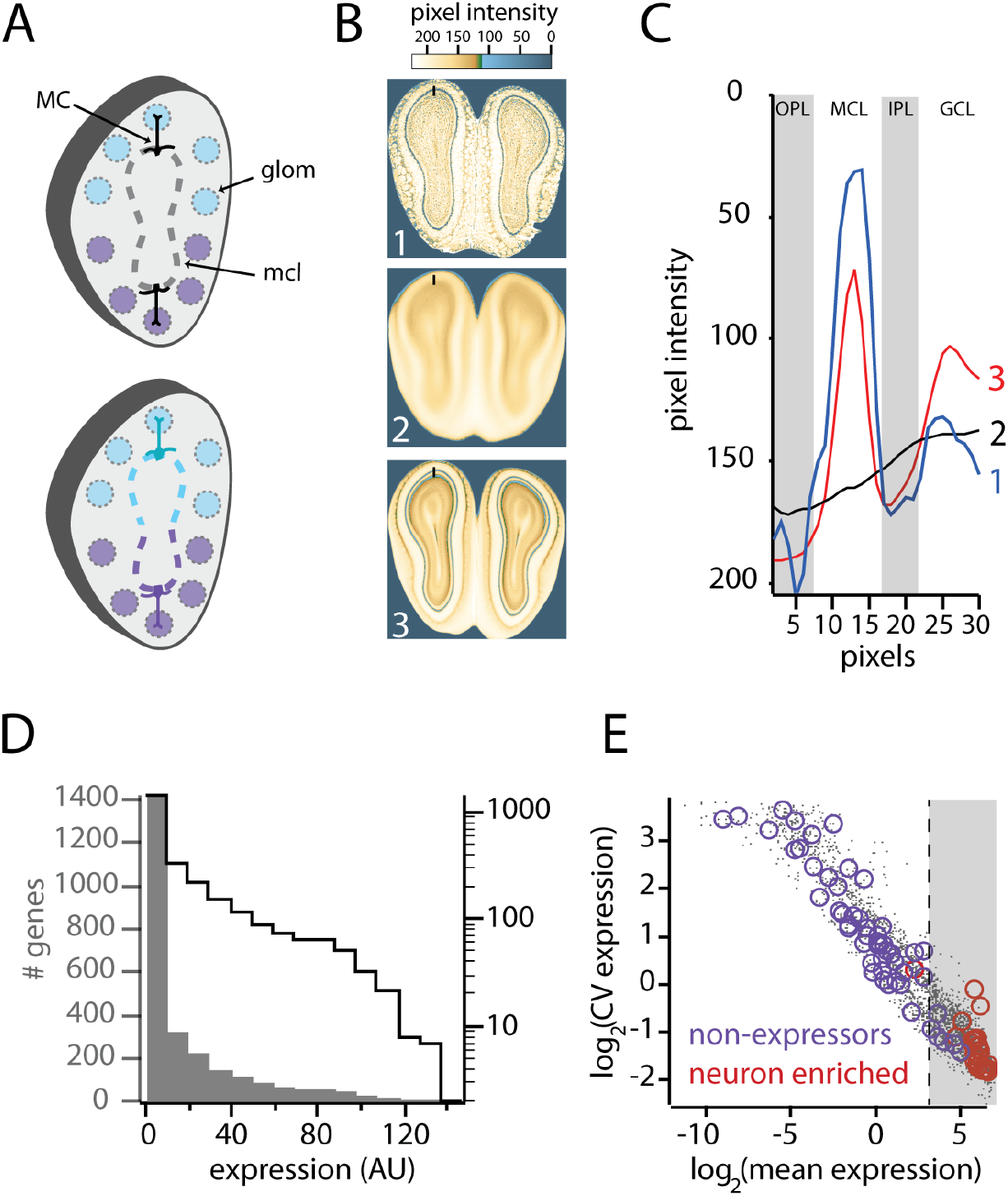
A genomic-scale atlas of registered in-situ-hybridization (ISH) experiments for the mouse olfactory bulb derived from the Allen Mouse Expression Atlas. **A)** Two contrasting models of mitral intra-laminar heterogeneity. *Top:* mitral cell population is homogeneous, providing invariant readout of zonally organized inputs. *Bottom:* mitral cell population is heterogeneous (comprised of cellular and/or local circuit-subtypes) for zonally organized readout and computation. **B)** Demonstration of successful large-scale image registration. *1)* example of a single registered ISH image; *2)* median intensity projection of ~2K pre-processed (unregistered) ISH images; *3)* median projection image of the same images, post-registration. Note that figures (**B**) and (**C**) are reproduced in part from a previous methods-paper (Andonian et. al, 2019). **C)** Line-intensity profiles, calculated across the lines of interest shown in **B1-3**, demonstrating clear delineation of the mitral cell layer in registered images. Abbreviations: *OPL* - outer plexiform layer; *MCL* - mitral cell layer; *IPL* - inner plexiform layer; *GCL* - granule cell layer. **D)** Histogram of median expression level of all genes in the mitral cell layer. In agreement with previous work, the majority of genes show little or no expression. Left axis (corresponding to filled gray bars) is linear, and right axis (corresponding to black line) is logarithmic. **E)** CV vs. mean of gene expression in the mitral cell layer, for all genes in this study (gray dots). Purple and maroon circles are genes previously identified as ‘non-expressors’ and ‘neuron enriched’, respectively, in Lein et al. 2007. Gray area shows genes retained for the present study, with the dashed line indicating the chosen expression threshold.

While zonal differences in the OB’s inputs and projections are well appreciated, it is not clear whether the OB’s second-order neurons (mitral cells, in particular) and their associated local circuits show matching zonal organization. At one extreme (Fig. 1A, top), the mitral cell layer (MCL) may comprise a homogeneous cell population that provides invariant readout across glomeruli. At the other extreme (Fig. 1A, bottom), mitral cells may be both heterogeneous and spatially organized for input-specific readout and formatting of sensory inputs. While other studies have observed heterogeneity in mitral cell physiology^10^, and described its organization at the single-glomerulus scale^11–13^, the critical question of whether mitral cells show zonally organized differences has not been directly addressed.

To distinguish between the models of intra-laminar heterogeneity above (Fig. 1), we first investigated spatial gene expression profiles along the mitral cell layer, at genomic scale (i.e. for thousands of genes). In contrast to the serial and piecewise examination of individual marker genes, this unbiased approach investigates the high-dimensional organization of transcription, in search of region-defining expression motifs. Similar approaches have been used in the hippocampus to confirm the intra and inter-laminar differences of classical histology^14,15^, as well as to discover new ones ^16^

We fetched all (~30,000) coronal in-situ-hybridization (ISH) experiments for the olfactory bulb from the Allen Institute’s mouse brain expression atlas^17^, and spatially registered these into a high-dimensional atlas, allowing clear differentiation of the OB’s laminae (Fig. 1B,C). Briefly, our registration pipeline used a convolutional neural network (CNN)^18^ to classify sections and design templates for non-affine registration of ISH experiments (See Methods; Supp. Video 1). Methods are described more thoroughly in a previous publication^19^. For ease of interpretation, our analysis only included relatively rostral sections of the OB, in which the MCL is closed and contiguous. Expression energy (see methods) was calculated along the mitral cell layer in 193 contiguous 2D (5 pixel x 5 pixel) bins.

Consistent with previous results^15,17^, the majority of genes showed either no expression or negligible expression in the mitral cell layer (Fig. 1D). Indeed, there was strong agreement on the identities of enriched and non-enriched genes between our curated data set and previously published work (Fig. 1E). To validate our extracted expression values against a second source using a different assay of gene expression, we compared median expression values in our mitral cell atlas with those from the publically available spatial transcriptomic RNA-seq set^20^(Fig. S1). Again, we observed a strong correlation between the two data sets (R=0.57, p<8.63 x 10^-22^).

Figure 2A shows the matrix of gene-expression x distance along the mitral cell layer which we analyzed for potential low-dimensional structure. Scanning across all genes (i.e. across rows of the matrix), we observed greater differential enrichment in comparisons between dorsal and ventral locations relative to comparisons between adjacent locations (Fig. 2B). As shown in figure 2C, the histogram of pairwise distances (that is, distances in *gene-space)* between all MCL points was multimodal, indicating that gene expression profiles are not uniform across this cell layer, and may instead aggregate into several discrete types. To further investigate this possibility, we performed a “component-based” clustering of expression profiles using non-negative matrix factorization (NMF^15,21^; see methods; Fig. S2). For a 2-d decomposition, we observed prominent dorsal and ventral spatial modes describing gene expression (Fig. 2D). Notably, these clustering results reflect strictly *genomic* distances between points along the mitral cell layer, and make no explicit use of spatial information in the clustering process. More plainly: the NMF results suggest that many individual genes show strong and selective dorsal or ventral patterning.

**Figure 2.**
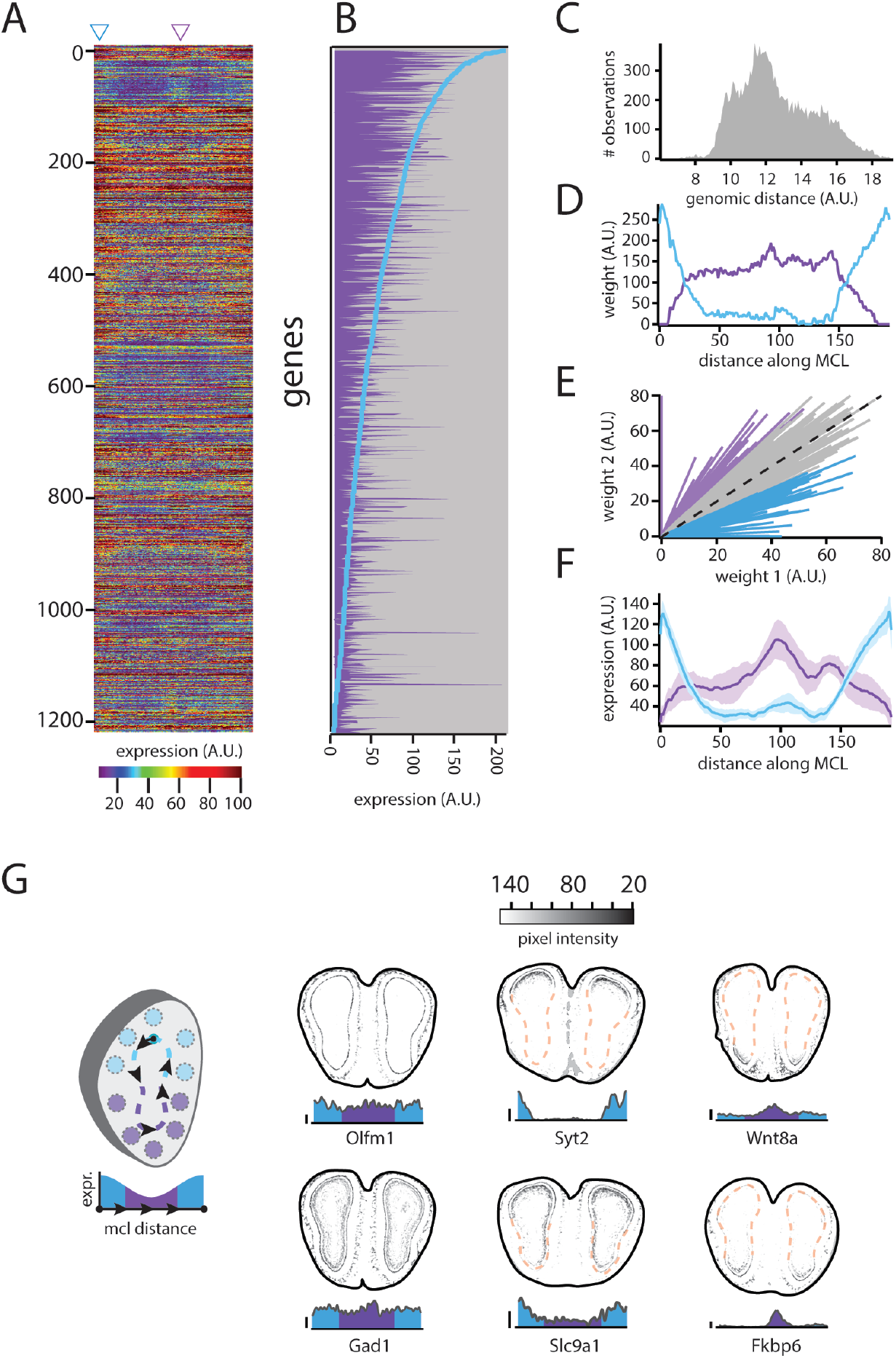
Clustering of mitral cell layer (MCL) gene expression profiles. **A)** Matrix of genes (rows) x distance along the MCL (columns). Left-most column is expression data from the most dorsal point of the mitral cell layer. Consecutive columns ‘trace’ the mitral cell layer counter-clockwise, until arriving at the same starting dorsal point (see schematic in panel **G**). **B)** Illustration of dorso-ventral gene enrichment. Blue line shows rank-ordered gene expression values from a dorsal location in the MCL (column indicated by teal triangle in (**A)**); Purple bar-plot shows gene expression values from a ventral location (purple triangle in (**A**)), ordered according to the rank-ordered dorsal values. Note that while the overall trend is similar, there are genes that are highly expressed ventrally that are virtually unexpressed dorsally. The converse is also true, but difficult to see in the densely packed bars. **C)** Histogram of pairwise distances (Euclidian distance in gene-expression space) between all pairs of 193 points along the mitral cell layer. **D)** Gene expression clusters (i.e. ‘features’) derived from performing non-negative matrix factorization (NMF) on the matrix in (**A**), showing prominent dorsal (teal) and ventral (purple) modes defining transcription in the MCL. **E)** Clustering of individual mitral cell gene expression profiles by their relative NMF weights (i.e. by their resemblance to the either mode 1 or mode 2 in (**C**)). Each radial line corresponds to one gene, with expression amplitude indicated by length, and degree of clustering indicated by angular deviation from the identity line (dashed line). Teal and purple lines are genes considered ‘strongly clustered’ dorsally and ventrally, respectively (strong clustering is defined here as those genes with angular deviations >1.5 SDs from the mean of the angular distribution). **F)** Averaged expression profiles for the 20 most strongly expressed dorsal and ventral genes. Shaded area is S.E.M. **G)** Sample expression profiles of individual genes. *Left schematic:* convention for plotting expression profiles. Profiles begin at the dorsal-most point of the MCL, and proceed counter-clockwise along the extent of the MCL. *Right panels:* examples of genes with strong uniform expression (left column); strong and selective dorsal expression (middle column); strong and selective ventral expression (right column). Raw & unadjusted ISH images corresponding to data in (**G**) are shown in Supplementary Figure S3)

Accordingly, expression profiles of individual genes were readily clustered by their resemblance to one of the two given modes (and to the exclusion of the other), and figure 2E shows that there are many candidate genes with strongly biased dorsal vs. ventral expression. In principle, it is possible that individual expression profiles do not show categorical dorsal or ventral enrichment, and that the derived components are instead a decomposition of a complex, multi-peaked expression profile shared by many genes. Although the histogram of Fig. 2C already argues against this, we investigated this further by averaging the expression profiles of the top 20 genes in the dorsal and ventral clusters. The averaged profiles strongly resemble the components derived from NMF (Fig. 2F), arguing that there exist categorical dorsal vs. ventral expressors.

Examples of individual gene expression profiles are shown figure 2G. The left column shows examples of strong uniform expressors (Olfactomedin 1 and Gad1), the middle column shows examples of strong dorsal expressors (Synaptotagmin 2 and Slc9A1) the right column shows examples of strong ventral expressors (Wnt8A, and Fkbp6). Raw images of these same ISH experiments are provided in Fig. S3, and argue against the possibility that asymmetries simply reflect pathological cases in which part of the mitral cell layer was missing or damaged. Taken together, our analysis indicates that many genes in the OB mitral cell layer exhibit dorsal and ventral patterning, in apparent correspondence with the dorsal and ventral patterning observed for the bulb’s sensory inputs.

While many of the genes showing spatially localized enrichment are involved in cellular excitability and synaptic transmission, it is still possible that this does not manifest as physiologically observable or meaningful differences in mitral cell properties. As in other systems, there may simply be different genetic programs capable of producing phenotypically equivalent mitral cells^22,23^. Additionally, it is possible that the observed transcriptional differences are only important for cellular migration and axonal targeting, and not excitable properties or local computation.

To explore this possibility, we made whole-cell recordings from coronal sections of the olfactory bulb, and compared physiological properties between dorsal vs. ventral mitral cells (see Methods). We observed a marked difference in the firing rate vs. input (FI) relationship for the two cell types, with ventral mitral cells exhibiting steeper FI gain. Peak firing rate for dorsal mitral cells was 42.85±7.03 Hz, whereas peak firing rate for ventral mitral cells was 79.90±12.19 Hz (p=0.02, unpaired t-test, n=15 in both cases). Input resistance and resting potential did not differ for the two cell populations (R_input_(dorsal)=113.26 ±19.62 vs. R_input_(ventral)=148.93±44.67, p=0.43, unpaired t-test, n=12 and n=11; (V_rest_(dorsal) = 55.2±1.9 mV vs. V_rest_(ventral)=57.1±2.3 mV, p=0.51, unpaired t-test, n=13 and n=15, respectively). On inspection of all individual FI curves (Fig. 3C), it was evident that the ventral population contained a subset of cells supporting higher firing rates; this is additionally suggested by the greater variance of peak firing rates in ventral mitral cells (Fig. 3C).

**Figure 3.**
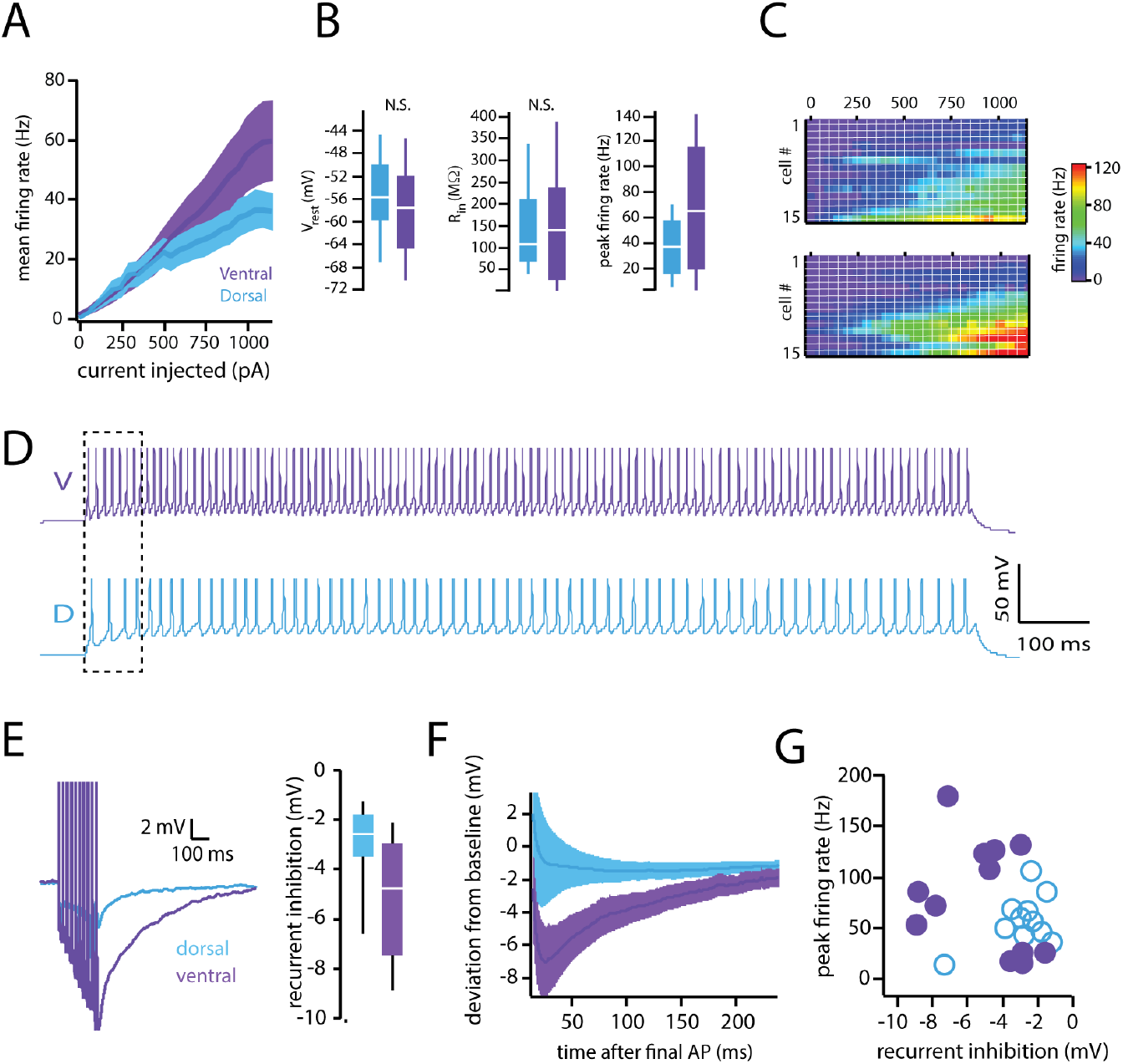
Electrophysiological differences between dorsal and ventral mitral cells. **A)** Action Potential Frequency vs. input (FI) relationship for mitral cells. Stimuli were 1*s* injections of step-currents in current clamp (see Methods). **B)** Dorsoventral differences in intrinsic properties of mitral cells. **C)** Montages summarizing all experiments investigating the FI relationship between dorsal and ventral mitral cells. **D)** Traces showing APs observed in dorsal and ventral mitral cells, in response to the stimulus evoking peak firing. **E)** *Left:* Traces showing recurrent inhibition in dorsal and ventral MCs elicited by a fixed number of supra-threshold stimuli. *Right:* Differences in peak amplitude of recurrent inhibition in dorsal and ventral mitral cells. **F)** Timecourse of recurrent inhibition for dorsal and ventral MCs. **G)** Scatterplot of firing rate and amplitude of inhibition, for all cells with both measurements.

We also tested for differences in the magnitude of recurrent inhibition in dorsal vs. ventral mitral cells by eliciting a fixed number of action potentials, and measuring the slow, AHP-like potential after the final spike. This potential is known to reflect local circuit dynamics-specifically, recurrent dendrodendritic inhibition mediated by granule cells^24–26^. We again observed a marked difference between dorsal and ventral mitral cells, with ventral cells having showing ~ 70% more inhibition than dorsal cells (Peak AHP amplitude (dorsal) = – 2.92±0.52 mV; Peak AHP amplitude (ventral)= −5.08±0.69 mV, p=0.02, unpaired t-test, n=12 and n=13, respectively). When peak MC firing and peak MC inhibition were plotted on the same axes, a subset of ventral cells were characterized by both high firing rates, and greater recurrent inhibition. Under the assumptions of a simple model in which mitral cells are presumed to merely gate throughput to downstream structures, these two effects appear to be in opposition. One possibility is that the more pronounced inhibition is a homeostatic adjustment to counter-act greater excitability. Of course, these differences are not necessarily opposed, functionally, and may play more complex roles in shaping local population dynamics (synchrony, oscillatory behavior, etc) as well as the spatiotemporal formatting of activity for downstream targets with different ecological roles^27^.

In sum, we combined ‘genomic-anatomical’ and physiological analyses to show that olfactory bulb mitral cells are heterogeneous and zonally organized. We note that our analysis only establishes a lower bound on this heterogeneity, and does not exclude the possibility that there are additional mitral cell or local circuit subtypes. One intriguing possibility raised by our study is that inputs to the dorsal vs. ventral bulb undergo distinct local formatting, in a manner that is either input or target specific (or both). A limitation of our study is that our methods cannot unambiguously differentiate between a model in which there are two, unmixed and non-overlapping populations of different mitral cells, versus two mixed populations that differ in relative abundance in the dorsal vs. ventral bulb. Future work employing single-cell sequencing techniques will be able to address this more directly.

## Methods

### Image registration

Our registration and image-processing methods - including comparisons to other methods, and analysis of registration accuracy -- are described in detail in a recent methods-paper from our group^17^. Code for all described procedures is available at https://github.com/CastroLab/ImageRegistrationPipeline. Briefly, our workflow consists of data fetching, pre-processing, classification and registration template design, registration, and clustering.

#### 1) Image fetching

All (~ 30,000) in-situ hybridization (ISH) coronal experiments containing olfactory bulb were downloaded from the Allen Brain Atlas (ABA), using the Allen Institute’s API and our own custom-written Python scripts. In addition, we also downloaded the expression image masks for each ISH experiment, which are thresholded and segmented image masks derived from the raw data that map expression differences to a common 8-bit scale. Images were JPEGs ~ 2MB each and were maintained on local servers. Details on the ABA tissue-processing pipeline have been described in detail in previous publications, and are also documented on publicly available white-papers from the Allen Institute: (http://help.brain-map.org/display/mousebrain/Documentation).

Each ABA coronal tissue section is annotated with a unique section-ID indicating rostro-caudal position relative to a reference brain sectioned at 100 μm intervals. We grabbed all tissue sections with section IDs > 400 (larger values are more rostral, and the bulb is the most rostral structure of the mouse brain). This conservative range was chosen empirically to miss no sections with olfactory bulb (OB). Owing to this conservative criterion, we initially retained some sections that did not contain OB, and which included portions of frontal cortex, piriform cortex, and the anterior olfactory nucleus (AON).

#### 2) Image pre-processing

Custom-written Python scripts were used for segmentation, thresholding, cropping, and affine transformations to correct for differences in image scale as well as slight differences in image orientation. Images were converted to grayscale, centered, and downsampled to 220 × 220. After thresholding, the binary segmentation mask underwent a morphological closing operation (skimage.morphology.binary_closing) with a disk structuring element of radius 2. This mask was used to set all background pixels to pure white, which was critical for registration.

#### 3) Derivation of a CNN-based image similarity metric

We trained a convolutional neural network (CNN) to classify OB tissue sections into one of 6 groups (5 anatomical section types & 1 ‘other’ category comprising damaged, low-quality, or otherwise un-classifiable sections). The network was trained on a random set of 2,227 images that were expert-labeled (by JBC). The classifier was implemented in MatConvNet (vlfeat.org)^28^ and we used a transfer learning approach using a pre-trained architecture (the popular Alexnet architecture^29^, trained on ImageNet (www.image-net.org)), and fine-tuned only the last fully connected layer and softmax classifier. Briefly, the architecture comprises 5 Convolutional/Rectified Linear Unit (ReLU) layers, three of which are followed by max pooling, and three fully connected (“dense”) layers, for a total of 7 hidden layers and an output layer. The network was trained on a custom-built machine with 2 x SLI Dual GeForce GTX Titan x 12GB GPUs, and training took ~2-3h. All downloaded ABAISH images were passed through the trained CNN, and the weights of the 7th (and final) fully-connected layer (fc7) were used as feature vectors to describe the images. Pairwise Euclidian distances between these fc7 vectors were used as a proxy for image similarity.

To design a set of registration templates, we chose 12 uniformly distributed seed-points across the image space of closed MCLs to define a grid. We then hand-picked 10 individual ISH experiments within each of these 12 neighborhoods, choosing images for symmetry, tissue quality, and contrast. These sets of ten images were registered groupwise using the Medical Image Registration Toolbox (MIRT)^30^-a Matlab-based package for performing parametric (B-spline) registrations. Registrations (both groupwise and pairwise) were performed using cubic B-splines for image deformation, and similarity between the fixed and target images was assessed by calculating material similarity and mutual information^31^ as these metrics showed the least sensitivity to differences in contrast. Registration was performed with mostly default parameters, with the exception of regularization weight, which we increased from the default 0.01 to 0.1 to prevent biologically unmeaningful image deformations. The image transformations were discovered for ISH images and also applied to the corresponding expression images for subsequent quantification and analysis of gene expression. To ensure that derived expression profiles reflected genuine spatial variation in expression, and not artifacts introduced by tissue damage, poor registration, or other sources of error, registration quality for all registered images was ground-truthed by hand (by JBC, and DP). Approximately 72% of images were judged to be successfully registered, which is in accord with work in other systems ^32^

### Formatting and clustering of expression data

Expression data from of each registered image was extracted as a 1-D contour that traced the mitral cell layer (left bulb), beginning at the dorsal pole of this cell layer, and continuing counterclockwise until joining the original dorsal seed point. The mitral cell layer was tiled with 193 non-overlapping 5 x 5 bins, and at each bin the median value was computed and reported as the ‘expression value.’ Note that this is a unit-less measure from 0-255 in which pixel intensity serves as a proxy for the intensity of gene expression. While there is no straightforward way to report this number as an RNA count or a FPKM we did observe a strong rank correlation between our extracted expression values, and those from the publicly available Spatial Transcriptomic (ST) database^33^ (data expressed as Transcripts per Million (TPM), see Fig. S1). ST expression data are uniformly sampled across coronal sections of the OB in spots of diameter 100μ, with center-to-center distances of 200μ between spots. We used the R-based package SpatialCPie (http://bioconductor.org/packages/devel/bioc/vignettes/SpatialCPie/inst/doc/SpatialCPie.html) to extract spot-data from the left mitral cell layers of all 12 available ST replicates. Only spots which spanned the MCL exclusively and unambiguously (i.e. including no portions of the granule cell layer) were chosen. This yielded data from a median of 3 spots per ST replicate. Median TPM was computed for all genes across all spots and all replicates to generate a super-set of ST expression values. The set of these genes concordant with the Allen genes we curated is what is reported in figure S1.

Non-negative matrix factorization (NMF) is a popular method of dimensionality reduction for genomic data, owing the intrinsic non-negativity of gene expression, and the ease of interpreting NMF-derived clustering results. NMF-based methods have been used in component-based analyses of gene expression in the hippocampus^15^, as well as in the olfactory bulb as a whole. Briefly, NMF seeks a low-dimensional approximation of an m (genes) x n (bins) data matrix, **A**, as the product of two lower dimensional matrices^21^: A matrix of features, **W** (m x s) and a matrix of weights, **H** (s x n). The new dimensionality, s, is chosen such that s<<m. We employed Matlab’s standard implementation of NMF (nnmf.m) using the alternating least-squares algorithm, and using a maximum of 1,000 iterations. The data matrix, **A**, was divided into random but equal-sized training and testing halves for each of 250 runs of the factorization. Because the factorization is not unique, we monitored the consistency of the derived basis vectors (**W_i_**) in consensus matrices (Fig. S2), and observed that clustering for s=2 was strong and reliable. Briefly, we initiated an m x m connectivity matrix, **C**, and updated its entries by 1 for each NMF run. That is, if genes i and j were co-clustered on a given run, then entry **Cij** was increased by 1. If the genes did not cocluster, entry Cij was unchanged. If clustering is stable and basis vectors are consistent, we expect that the distribution of elements of C after all runs will be strongly bimodal, with entries aggregating at 0 and 1, as in figure S2. We reported the most stable version of the basis vectors by computing KL divergence between all pairs of the 250 instances of W, and selecting W with the lowest mean KL divergence value.

#### Electrophysiological methods

Acute coronal OB slices (300μ) were prepared from P21-P28 C57/Bl6J mice (Jackson labs) using a vibratome (Campden Instruments). Mice were anesthetized with isoflurane, decapitated, and their brains removed and submerged in icecold oxygenated (95/5 O_2_/CO_2_) artificial cerebrospinal fluid (ACSF) with the following composition (in mM): 125 NaCl, 2.5 KCl, 25 NaHCO_3_, 1.25 NaH_2_PO_4_, 1MgCl_2_, 25 glucose, 2 CaCl_2_. Following sectioning, slices were incubated in the same oxygenated ACSF at 37°C for at least 45 minutes before beginning physiology experiments. Only sections with a contiguous MOB mitral cell layer (uninterrupted by the accessory olfactory bulb) were used for experiments. After incubation, slices were transferred to an electrophysiology rig, where they were perfused with the same oxygenated ACSF solution, and visualized under DIC optics (Olympus BX51). Whole cell recordings were obtained with glass electrodes (2-8MΩ) filled with the following internal solution (in mM): 150 potassium gluconate, 2 KCl, 10 HEPES, 10 sodium phosphocreatine, 4 MgATP, and 0.3 Na3GTP, adjusted to pH 7.3 with KOH. Recordings were made using a Multiclamp 700B amplifier (Molecular Devices), and digitized at 10kHz with an ITC-18 A/D board. Data collection was performed using custom-written routines in Igor Pro (Wavemetrics). When the slice was first placed in the recording chamber, x,y coordinates of the micromanipulator (Sutter) were marked for the most dorsal and most ventral points along the mitral cell layer. To be included as a “dorsal” cell for subsequent analysis, the y coordinates of a given cell body had to be in the dorsal-most third of the mitral cell layer (as defined by the previously measured extreme points). Similarly, to be included as a “ventral” cell for analysis, the cell body had to be in the ventral-most third most third of the mitral cell layer. All experiments (measurements of input resistance, firing rate, and recurrent inhibition) were performed in current clamp, with the experimenter manually adjusting holding current in between sweeps to maintain cells near −55 mV. The only exception to this was on sweeps when resting potential was explicitly measured, and no holding current or stimulus was applied. Cells with high spontaneous activity, or for which this target voltage could not be maintained with < 150 pA current injection were not included for analysis.

Stimuli used included the following: 1) Hyperpolarizing steps (to measure input resistance) of 1s duration, from −50pA to −200pA, in 4 steps of 50 pA; 5 blocks for a total of 20 sweeps. 2) Depolarizing steps (to measure the FI relationship) of 1s duration, from + 50 pA to +1200 pA, and proceeding in steps of 50 pA for 24 sweeps (3 blocks total); 3) Supratheshold current pulses (to measure recurrent inhibition): 10 pulses at 40Hz, each pulse 2ms in duration; 30 total sweeps. Starting pulse amplitude was 2nA, and was increased if necessary to evoke supratheshold responses to all 10 pulses in a sweep. Sweeps in which APs did not occur for all 10 stimuli were not included in the analysis. Data analysis was performed using custom-written routines in IgorPro (Wavemetrics) and Matlab (MathWorks), and was done with analysts blinded to cell identity (dorsal vs. ventral). Except where otherwise indicated, all electrophysiological measurements are expressed as mean ± SEM. Box-plots in Figure 3 show group medians and 25^th^ and 75^th^ percentiles, with whiskers indicating 10^th^ and 90^th^ percentiles. Statistical significance of between-group comparisons was assessed using unpaired t-tests.

## Acknowledgements

We are grateful to the Allen Brain Institute for making their data publicly available. We thank Shreejoy Tripathy and Rick Gerkin for advice on data analysis, and Tina Rioux for technical assistance. This work was funded by an NSF CAREER grant (1553279) to JBC, and by an Institutional Development Award (IDeA) from the National Institute of General Medical Sciences of the National Institutes of Health under grant number P20GM0103423 to TJG.

## Author contributions

JBC conceived of and designed the study. DP and AA developed the CNN-based pipeline for image registration, with input and guidance from JBC. TG provided expertise on and assisted with image analysis. HL collected the electrophysiological data, with guidance from JBC. DP, AA, HL, and JBC analyzed the data, and prepared figures. AL developed the initial pilot-version of this work. JBC wrote the manuscript, with input from all other authors.

## Competing interests

The authors declare that they have no competing interests.

## Supplemental information

Included with this submission are:

1 Supplemental video

3 Supplemental figures

