## Supplemental figures and legends for "Spatially organized genomic and physiological heterogeneity of the olfactory bulb mitral cell layer"

**Supplementary information for Paseltiner et al.**

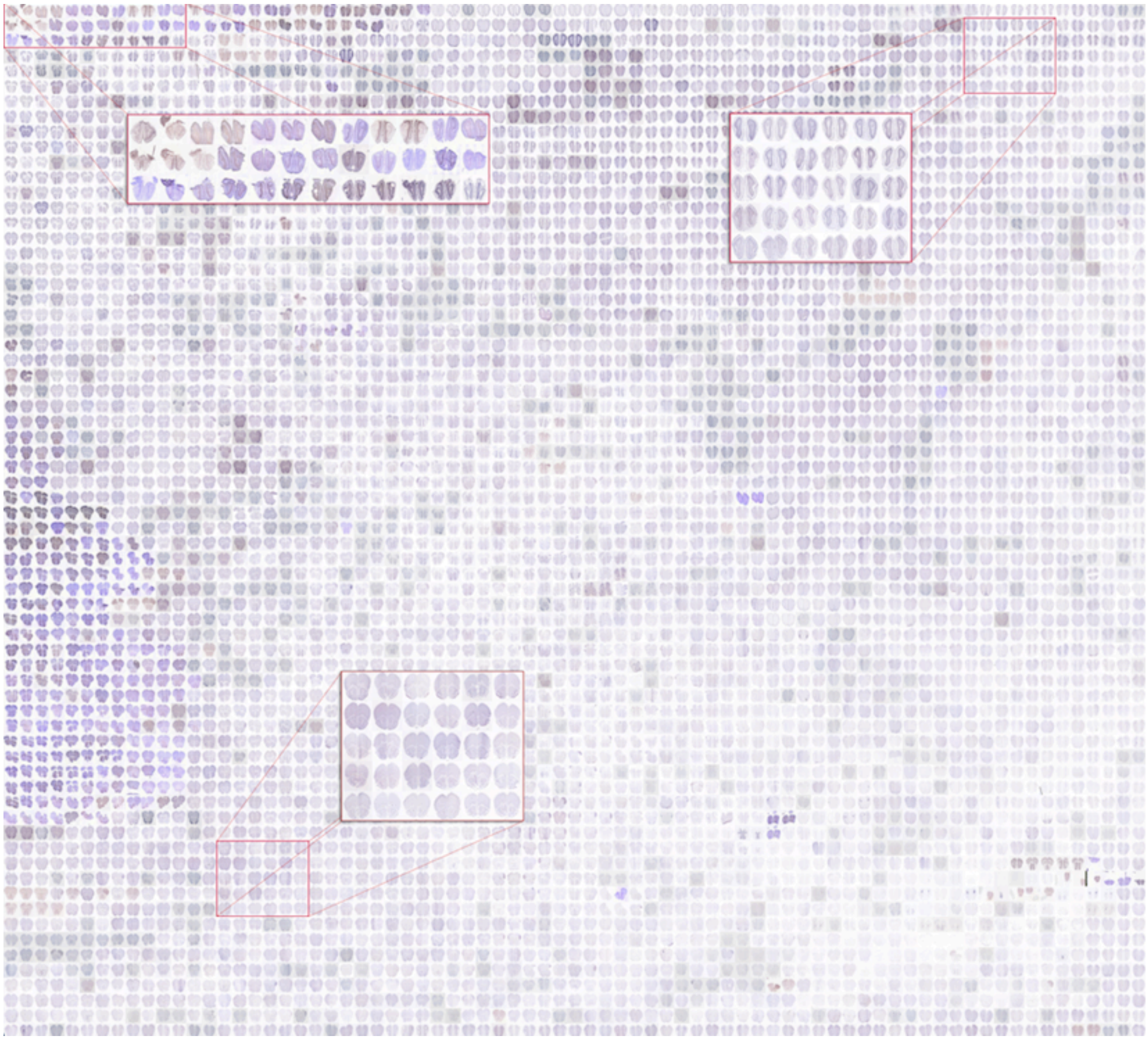

**Movie 1. A two dimensional embedding of ISH experiments from the Allen Brain Atlas.** Neighbor relationships between images were calculated from their respective fc7 vectors (see **Methods**, and Andonian et. al 2019[1]). Video begins with a panoramic view of a portion of image-space (~ 5,000 ISH experiments), and zooms into two different neighborhoods. Images shown are pre-processed but not registered.

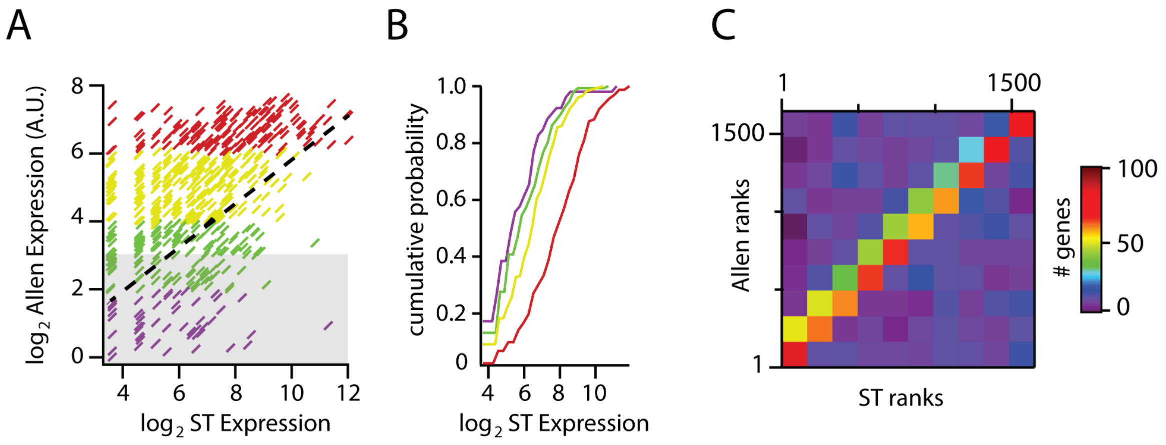


**Fig. S1. Agreement with Spatial Transcriptomic mitral cell expression data. A)** Plot of median mitral cell expression from Allen atlas vs. median mitral cell expression from the Spatial Transcriptomic data[2]. Dotted line is best least-squares linear fit (R=0.57, $p<8.63 \times 10^{-22}$). Data have been divided into quartiles (indicated by color) on the basis of their median (logarithmic) Allen expression levels, calculated for the MCL. Note that Allen expression values saturate at 256 (expression mask images are 8-bit (see methods)). Gray box indicates genes below threshold for inclusion. **B)** Normalized cumulative histograms of the data from A. **C)** 2D histogram (bin size=150 genes) of Allen Expression ranks and ST expression ranks, showing strong correspondence of ranks between the two data sets.

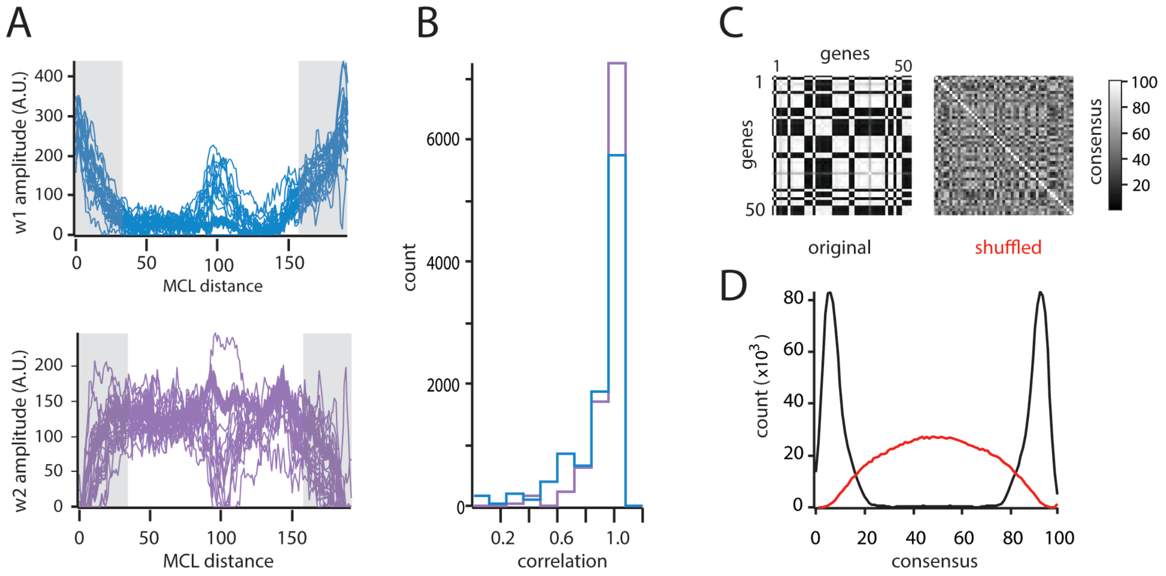


**Fig. S2. Consistency of NMF features and clusters. A)** Basis (feature) vectors computed for all 250 runs of the factorization. **B)** Histograms of correlation coefficients between all basis-vector pairs (computed separately for the dorsal (teal) and ventral (purple) biased vectors). **C)** Consensus matrix across NMF runs. For ease of viewing, only the first 50 genes are shown. Left is original data, and right is shuffled data. The tendency of pixels to be saturated black or white in the original indicates a high degree of consistency (i.e. gene pairs tend to either always co-cluster (black), or never co-cluster (white), with comparatively few genes having an ambiguous dorsal vs. ventral bias (intermediate gray values)). **D)** Histogram showing the distribution of consensus values. Data in (D) are from the entire data set, not just the subset shown in figure C.

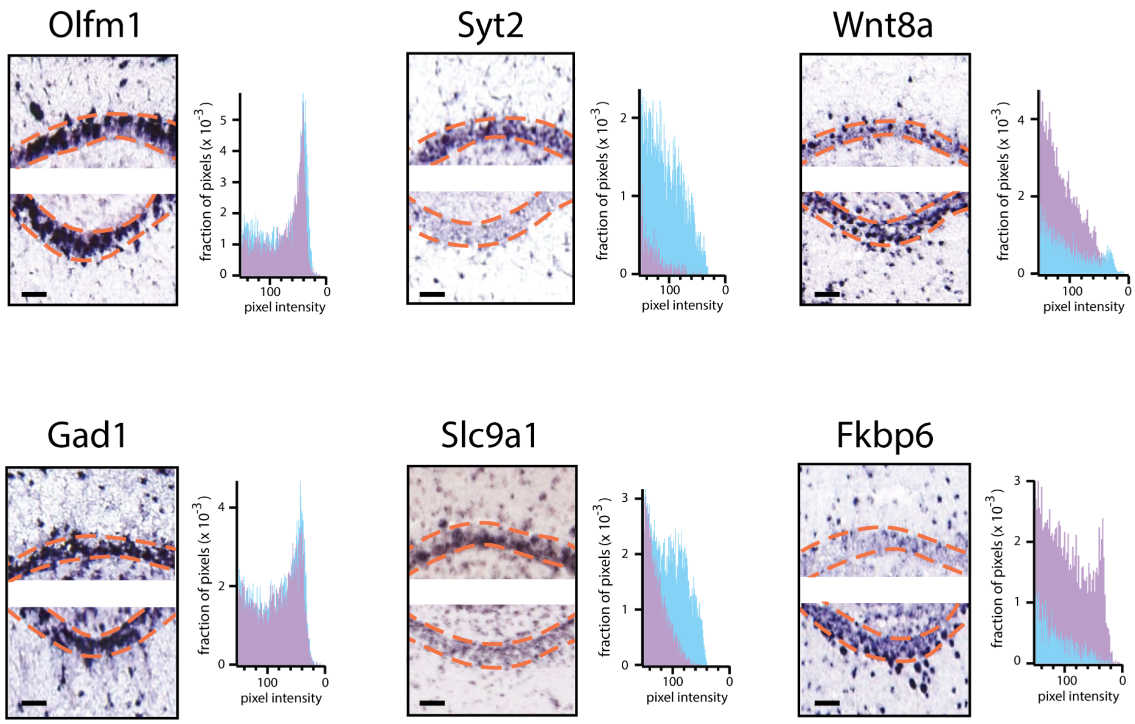


**Fig. S3. Raw, unadjusted in-situ hybridization data showing dorsal and ventral enrichment of individual genes in the mitral cell layer.** Data correspond to representations of expression shown in figure 2. Left column shows examples of uniform expression; middle and right columns show examples of dorsal and ventral enrichment, respectively. Insets are histograms of pixel intensity calculated for the dorsal (teal, top sub-panel of all images) and ventral (purple, bottom sub-panel of all images) mitral cell layers. Scale bar, 50μ in all images.
